# Energy Penalties Enhance Flexible Receptor Docking in a Model Cavity

**DOI:** 10.1101/2021.04.20.440636

**Authors:** Anna S. Kamenik, Isha Singh, Parnian Lak, Trent E. Balius, Klaus R. Liedl, Brian K. Shoichet

**Author notes:** These authors contributed equally (Anna S. Kamenik, Isha Singh). Corresponding authors. Trent E. Balius, Klaus R. Liedl, 0043-0512-50757100, Brian K. Shoichet, 01-415-541-4126. Author Contributions: ASK and IS equally contributed to the manuscript. A.S.K., I.S., T.E.B., K.R.L, and B.K.S. designed research; A.S.K., I.S., P.L. and T.E.B. performed research and analyzed data; A.S.K., I.S., T.E.B., K.R.L, and B.K.S. wrote the manuscript.

## Abstract

Protein flexibility remains a major challenge in library docking due to difficulties in sampling conformational ensembles with accurate probabilities. Here we use the model cavity site of T4 Lysozyme L99A to test flexible receptor docking with energy penalties from molecular dynamics (MD) simulations. Crystallography with larger and smaller ligands indicates that this cavity can adopt three major conformations, open, intermediate, and closed. Since smaller ligands typically bind better to the cavity site, we anticipate an energy penalty for cavity opening. To estimate its magnitude, we calculate conformational preferences from MD simulations. We find that including a penalty term is essential for retrospective ligand enrichment, otherwise high-energy states dominate the docking. We then prospectively docked a library of over 900,000 compounds for new molecules binding to each conformational state. Absent a penalty term, the open conformation dominated the docking results; inclusion of this term led to a balanced sampling of ligands against each state. High ranked molecules were experimentally tested by T_m_-upshift and X-ray crystallography. From 33 selected molecules, we identified 18 new ligands and determined 13 crystal structures. Most interesting were those bound to the open cavity, where the buried site opens to bulk solvent. Here, highly unusual ligands for this cavity had been predicted, including large ligands with polar tails; these were confirmed both by binding and by crystallography. In docking, incorporating protein flexibility with thermodynamic weightings may thus access new ligand chemotypes. The MD approach to accessing and, crucially, weighting such alternative states may find general applicability.

**Significance Statement:** The dynamic nature of biomolecules is typically neglected in docking screens for ligand discovery. Key to benefitting from various receptor conformations is not only structural but also thermodynamic information. Here we test a general approach that uses conformational preferences from enhanced and conventional MD simulations to account for the cost of transitions to high energy states. Including this information as a conformational penalty term in a docking scoring function, we perform retrospective and prospective screens and experimentally confirm novel ligands with T_m_-upshift and X-ray crystallography.

## Introduction

Proteins interchange between conformational states of varying probabilities (1). These rearrangements naturally also alter its physicochemical properties (2, 3). Exploiting these varying features can benefit ligand discovery (4, 5), but also presents several challenges. Key among them is weighting the different states by their energies, which has been shown to be crucial for docking success (4, 6); without such weights, high energy protein conformations, often better suited to ligand complementarity but harder to access, can dominate docking results, acting effectively as decoy conformations.

Structural models of proteins in distinct conformational states can be obtained from experiments like X-ray crystallography, NMR, or Cryo-EM. The choice of the single structure used for a docking campaign contributes to its likelihood of success, and choosing any single conformation inevitably leads to false-negatives even in successful campaigns. A solution to this problem is to consider multiple protein conformations, often referred to as ensemble docking or flexible receptor docking (4, 7–12). Yet, incorporating multiple protein conformations only increases accuracy in ligand discovery when they are weighted according to their ensemble probabilities (13, 14). When such energies have been incorporated in docking campaigns, they have enabled the discovery of ligands that are inaccessible to single state docking, often with high fidelity to subsequent structure determination of ligand-protein complexes. However, incorporating these weights has relied on experimental observables, such as occupancies from high-resolution structures. This has both limited the range of states that may be used—since states higher in energy than a few kcal/mol above the ground state will not be observed experimentally—and cannot be generalized to the vast number of targets for which such information is unavailable. Even when alternative conformational states can be observed in complex with different ligands (4, 9, 15, 16), their thermodynamic weights in the apo ensemble are typically unknown. It would be useful to have a general method of sampling and energy-weighting conformational states that would enable their exploitation in ligand discovery in general, and for molecular docking in particular.

In principle, computationally modelled protein structures, such as those derived from molecular dynamics (MD) simulations, can sample such conformational states (7, 17, 18), and can estimate their thermodynamic weighting (19, 20). In practice, challenges with many MD simulations include insufficient sampling, and the difficulty in weighting states by relative energies. The free energy minima, representing conformational states of a protein, are often separated by high-energy barriers, which are rarely overcome on time scales covered by conventional MD (cMD) simulations (1). Enhanced sampling algorithms, such as accelerated MD (aMD) (21), introduce a bias potential to lower the barriers between individual conformational states. This makes sampling of a diverse conformational ensemble, including higher energy conformational states, more efficient (22–24). A core question is whether the assumptions and approximations made in accelerated MD affect its ability to usefully weight the conformations sampled.

Here, we test energetic weights from MD simulation for ligand discovery in the engineered cavity site of T4 Lysozyme L99A (L99A). This hydrophobic cavity, first introduced by Eriksson, Morton, Baase and Matthews (25–28) as a model system to explore ligand binding and thermodynamics, has important advantages for exploring terms in ligand binding and docking-based discovery (in this study, protein flexibility). The cavity site is relatively small, only 150 Å^3^ in its apo state, and is completely enclosed from solvent in that conformation (**Figure 1A**). Combined with its dominance by apolar interactions, this simplifies the determinants of ligand binding. Despite its small size, there are still many hundreds of likely ligands that are readily available and testable from within docking libraries, enabling prospective predictions to test new docking terms and methods (9, 26, 27, 29–31). Previous studies have revealed at least 68 ligands for this cavity, many of which have protein-bound crystal structures determined (25, 32), enabling detailed retrospective studies. Despite its simplicity, L99A has complexities that make it interesting and relevant as a model site, and its thermodynamics (28, 33–35), dynamics (22, 36–44) and ligand (un)binding (27, 30, 45–53) have been intensely studied.

**Figure 1.**
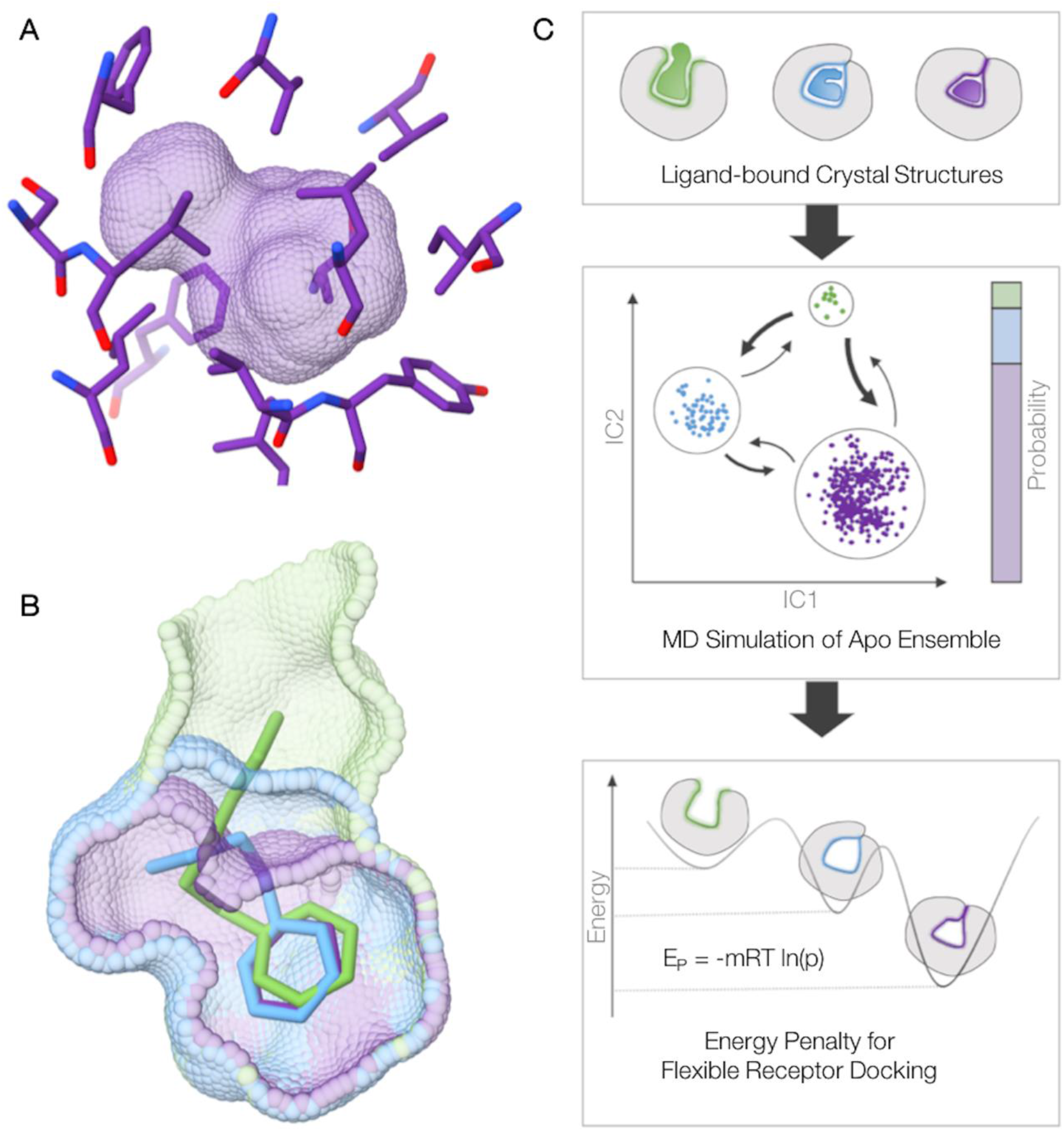
Three conformations of the L99A cavity binding site. **(A)** Crystal structures of T4 Lysozyme L99A in its apo state show a small, buried and entirely apolar cavity. **(B)** Structures of the protein in complex with ligands of increasing size show three major conformational states of the binding site: closed (purple), intermediate (blue) and open (green). **(C)** Workflow.

Particularly germane to this study, the cavity undergoes a conformational change as larger and larger ligands bind to it, adopting three principal conformations termed closed (150 Å^3^), intermediate (~200 Å^3^), and open (< 300 Å^3^) (29) (**Figure 1B**). As larger ligands bind, the cavity opens owing to the unwinding of helix F from an α- to a 3-10-helix. In the most voluminous state of the cavity (twice that of the closed state), it opens to form a channel between bulk solvent and the hydrophobic cavity. Thus, despite being a simplified model system, L99A exhibits substantial structural rearrangements, making it a useful site to test flexible receptor docking (9, 54).

For this study, we derive conformational state definitions from apo and holo crystal structures of L99A in its closed, intermediate and open state (**Figure 1C**). Removing the ligands, we perform aMD and cMD simulations for exhaustive and efficient sampling. We then construct a Markov state model (MSM) (55–58) to estimate the relative probability of each crystallographic conformational state in the apo ensemble. Converted into a conformational energy penalty *E_p_* (**equation 1**), we incorporate the state probabilities into our flexible receptor docking scoring function (4).

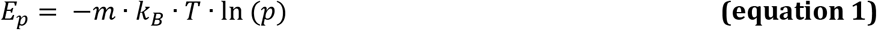

Where k_B_ = Boltzmann constant, T = temperature in K, p = population, m = weighting multiplier.

We test the reliability of the applied penalties in retrospective screens based on the known ligands and their property matched decoys. Crucially, we evaluate the ability of the approach to predict ligands with new chemotypes selective for each of the three relevant conformations of the cavity, in a prospective docking screen. We consider the usefulness of this approach to the general problem of predicting weighted conformational ensembles of proteins for docking and ligand discovery.

## Results

### Thermodynamic Weighting from Apo MD Simulations

We perform both enhanced, i.e., accelerated MD (aMD) (**SI Figure S1**), and conventional (cMD) simulations to estimate the population of the three crystallographic states in the apo ensemble (**Figure 2**). On the one hand, we directly derive reweighted populations from clustering 500 ns of aMD. This analysis suggests a probability of 0.5 % for the open, 1.4 % for the intermediate, and 98 % for the closed state (**SI Table S2**). On the other hand, we assess the stability of these populations from unbiased cMD simulations. From extensive structural sampling (7.75 μs) we construct an MSM to estimate the thermodynamic weight of each crystallographic state in solution with no ligand present. In addition to providing populations of and by extension weights for the states, this analysis also allows an assessment of the involved kinetics. The MSM analysis finds four distinct states (S_1_ to S_4_), which closely resemble the three distinct conformations observed in ligand-bound structures (**Figure 2, SI Figure S2–S4**). States S_1_ (0.05% of the population) and S_2_ (0.11% of the population) both resemble the open state, which, for example, occurs in complex with n-hexyl-benzene (PDB: 4W59). State S_3_ (1.09% of the population) resembles the intermediate conformational state, as found in the complex with n-butyl-benzene (PDB: 4W56). The closed conformation, which is found for most apo structures or in the complex with benzene (PDB: 4W52), is modelled by state S_4_ (98.76% of the population). Encouragingly, we thus obtain state populations within the same order of magnitude from the reweighted aMD ensemble and the unbiased simulations.

**Figure 2.**
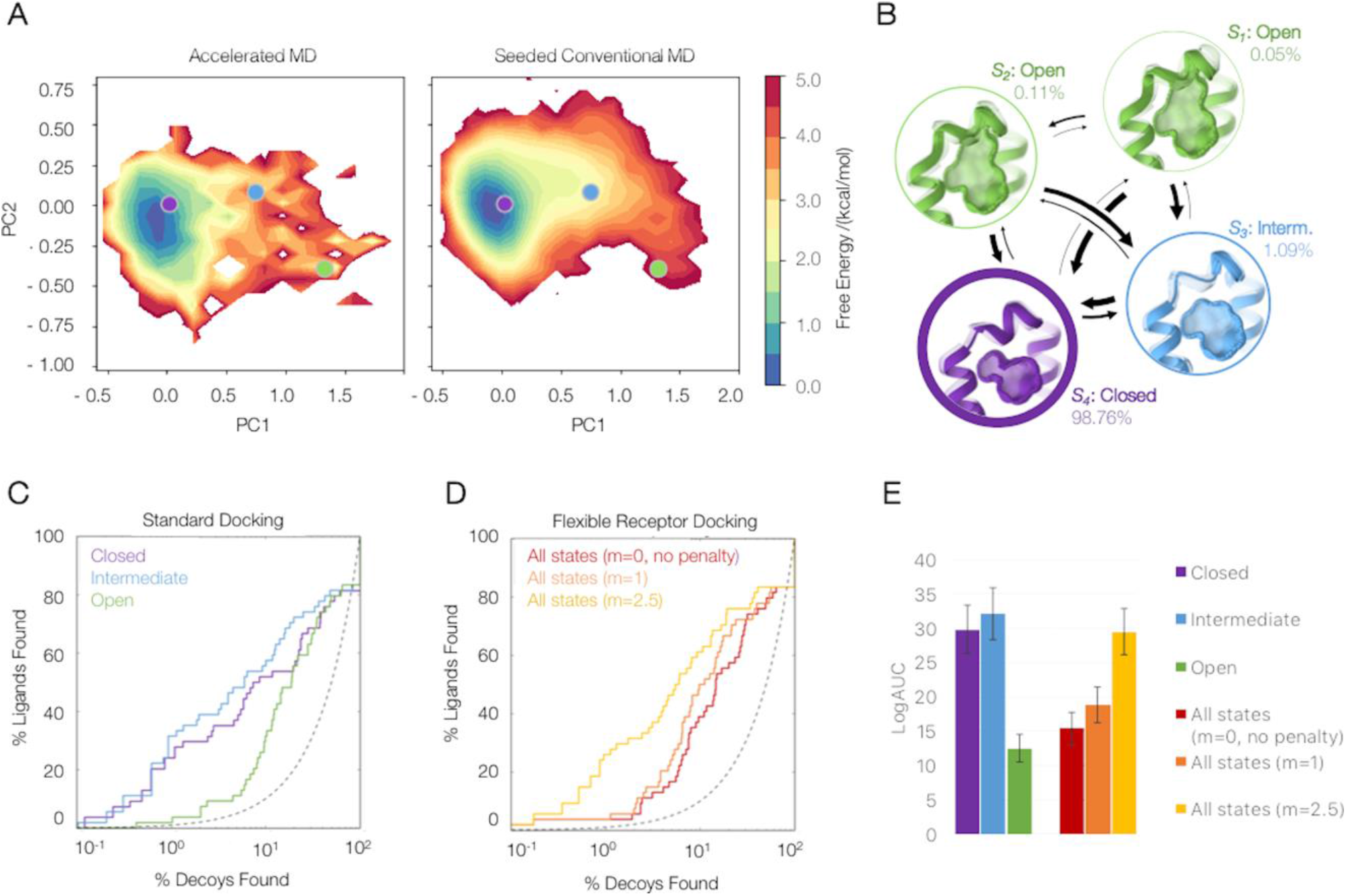
Retrospective docking with MD-based weightings. **(A)** Conformational space covered with aMD and the seeded cMD simulations as a projection onto the combined PCA space; Representative crystal structures of the closed (4W52, purple), intermediate (4W55, blue) and open (4W59, green) state are depicted as circles. **(B)** The MSM built from the unbiased cMD trajectories identifies four states (S_1_ to S_4_); S_1_ and S_2_ both represent the open state, S_3_ the intermediate and S_4_ the closed state of L99A. Adjusted log ROC curves from **(C)** standard docking to individual conformational states (purple: closed, blue: intermediate, green: open) and **(D)** flexible receptor docking with varying weighting multipliers m. **(E)** Adjusted areas under the log ROC curves quantifying enrichment of known ligands against decoys for classic and flexible docking screens with and without penalties.

The kinetic information provided by the MSM is not required to perform flexible receptor docking calculations. Nevertheless, this analysis highlights that the structurally highly similar closed and intermediate state have fast transition time scales, consistent with experimental spectroscopy (51, 59), and the ability to soak ligands into apparently closed cavities in crystals. Expanding the closed conformation toward the intermediate state is observed with a mean first passage time (mfpt) of 106 ns. Observing a full opening of the cavity, however, takes an order of magnitude longer in simulation time, with mfpts of 1555 to 8474 ns.

### Retrospective Testing of MD-based Penalties

We converted these populations into an energy penalty term using **equation 1** (4). To test the benefits and shortcoming of flexible-receptor docking and MD-based penalties, we docked 68 known ligands into the three conformational states of the binding site. We then calculated the log-adjusted enrichment of the known ligand over property matched decoys (60), docking to each receptor state individually (standard dock – **Figure 2C, 2E**) and using all three states combined with and without the energy penalty (flexible receptor docking - **Figure 2D, 2E**). As previously (4, 61), we find that enrichment of known ligands over decoys depends on the receptor conformational state. Docking to the higher-energy open state without a conformational penalty enables many decoy molecules to be accommodated, and moreover to score better than known ligands that bind to the smaller closed and intermediate states. Accordingly, docking against all three states without an energy penalty (**Figure 2D**, red curve) led to poor overall enrichment of known ligands over property-matched decoys and domination by the higher-energy open state (**Figure 2E**). Including a penalty term calculated directly from the MD slightly improves enrichment in the retrospective docking. When we weight this MD penalty term to bring it into line with the magnitude of the other energy terms in the DOCK3.7 scoring function, as similarly done in previous studies (4), ligand enrichment improves substantially, and the distribution of molecules docked to each conformational state becomes more balanced (less dominated by the open conformation).

In addition to its accuracy in identifying known ligands, we test whether the approach can reproduce the crystallographic loop state occupancies. We calculate the loop state propensities from the docking score of seven ligands as described previously (4). Applying a weighting multiplier of m=2.5 to the energy penalty term we find a striking Pearson correlation coefficient of 0.93 between experimental and predicted loop propensities (**SI Figure S5**).

### Prospective Screening to Identify New Binders for Each Conformational State

Encouraged by the retrospective performance, we applied the energy penalties in a prospective screen of a library of over 900,000 purchasable molecules from the ZINC database. From among the top-ranking 0.25% of these docked molecules, we selected 33 molecules for experimental testing, choosing the same number of compounds for each conformational state of the cavity (closed, intermediate, and open). Many of these molecules were topologically dissimilar to the previous known 68 ligands, with Tanimoto coefficient (Tc) similarities as low as 0.16 to any of the previous knowns (**Table 1**). Each of these was measured for binding by temperature of melting upshift using a Sypro orange binding assay (Methods). Of these 33, 10 increased the T_m_ of melting of L99A by between 1 and 5.5 °C when tested at 100 μM (p values ranging from <0.01 to 0.0001 vs. DMSO control) (**Figure 3**). In addition to the change in melting temperatures, we also determined crystal structures of 13 compounds bound to L99A. Taking together both change in melting temperature and crystal structures, we consider these 18 molecules to be L99A ligands, a docking hit rate of 55% (18 confirmed of 33 predicted, **Table 1**). On a state-by-state basis, 8 of 11, where the closed conformation was predicted by docking were confirmed experimentally, 7 of 11, where the intermediate conformation was predicted by docking were confirmed experimentally, while for the open state, which is more challenging because so many more ligands can fit it, 3 of 11 were confirmed to bind (**Figure 4**).

**Figure 3.**
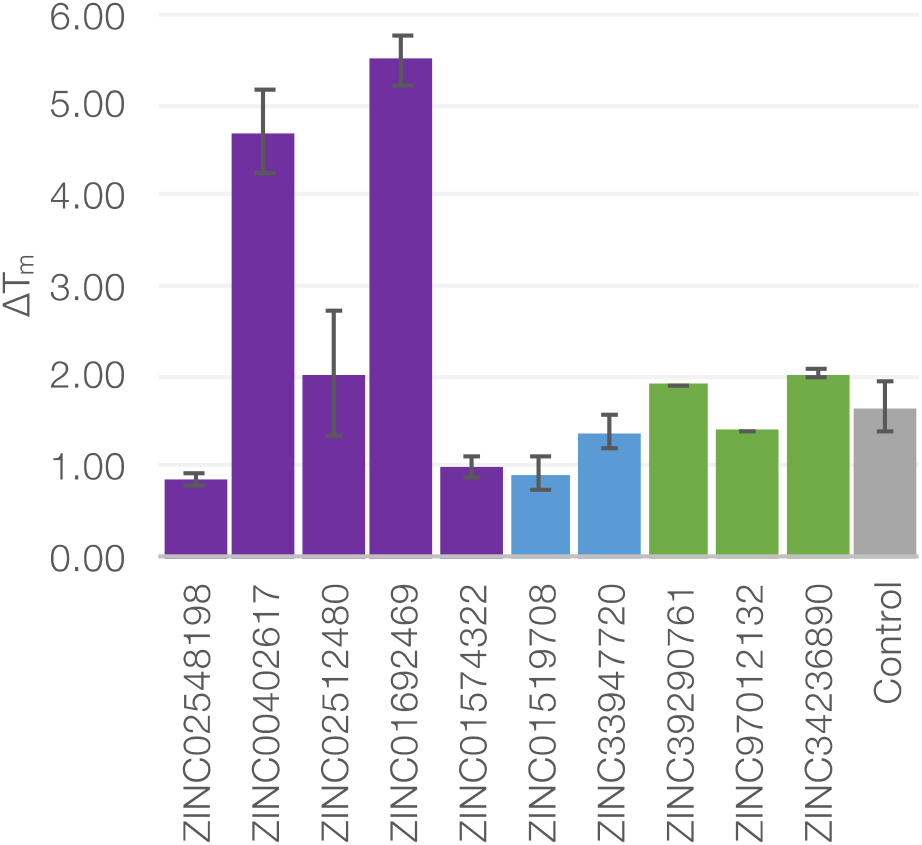
T_m_-upshift experiments with differential scanning fluorometry (DSF) In a thermal shift assay (DSF) twelve ligands were identified as binders. Five of which were predicted to bind to the closed state (purple), two to the intermediate state (blue) and three to the open state (green). The previously known binder Ethyl-Benzene is shown as positive control in gray.

**Figure 4.**
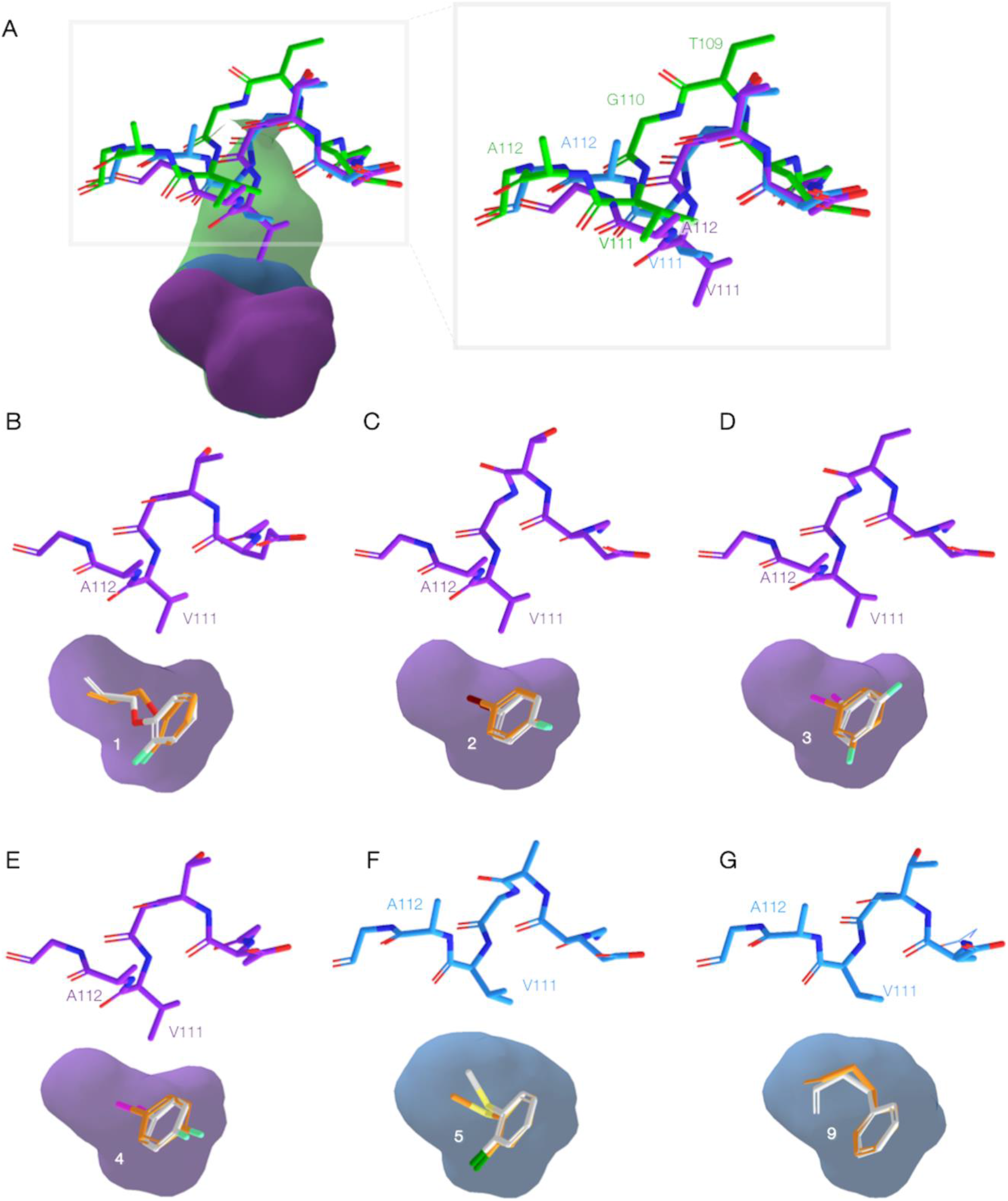
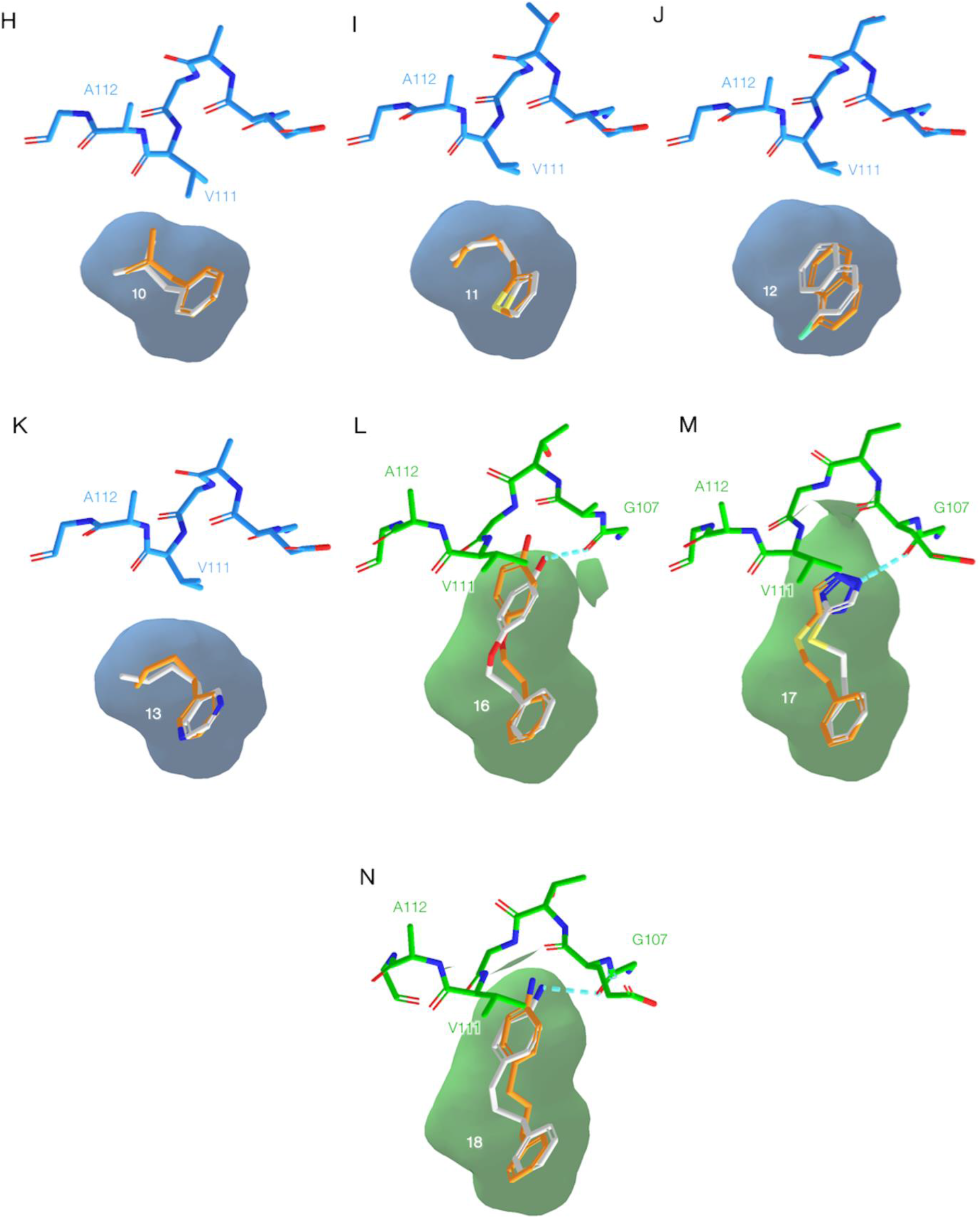
Predicted and experimental ligand poses and site conformations. **(A)** Closed (violet), intermediate (blue) and open (green) conformation of the L99A cavity site – characterized by a shift of the F-helix residues T109 to A112. **(B-N)** Superposition of predicted (orange) and crystallographic (white) ligand poses for crystals structures 7LOB, 7LOC, 7LOA, 7LOD, 7LX8, 7LX9, 7LOG, 7LOF, 7LOE, 7LXA, 7LX7, 7LX6 and 7LOJ respectively.

**Table 1.**
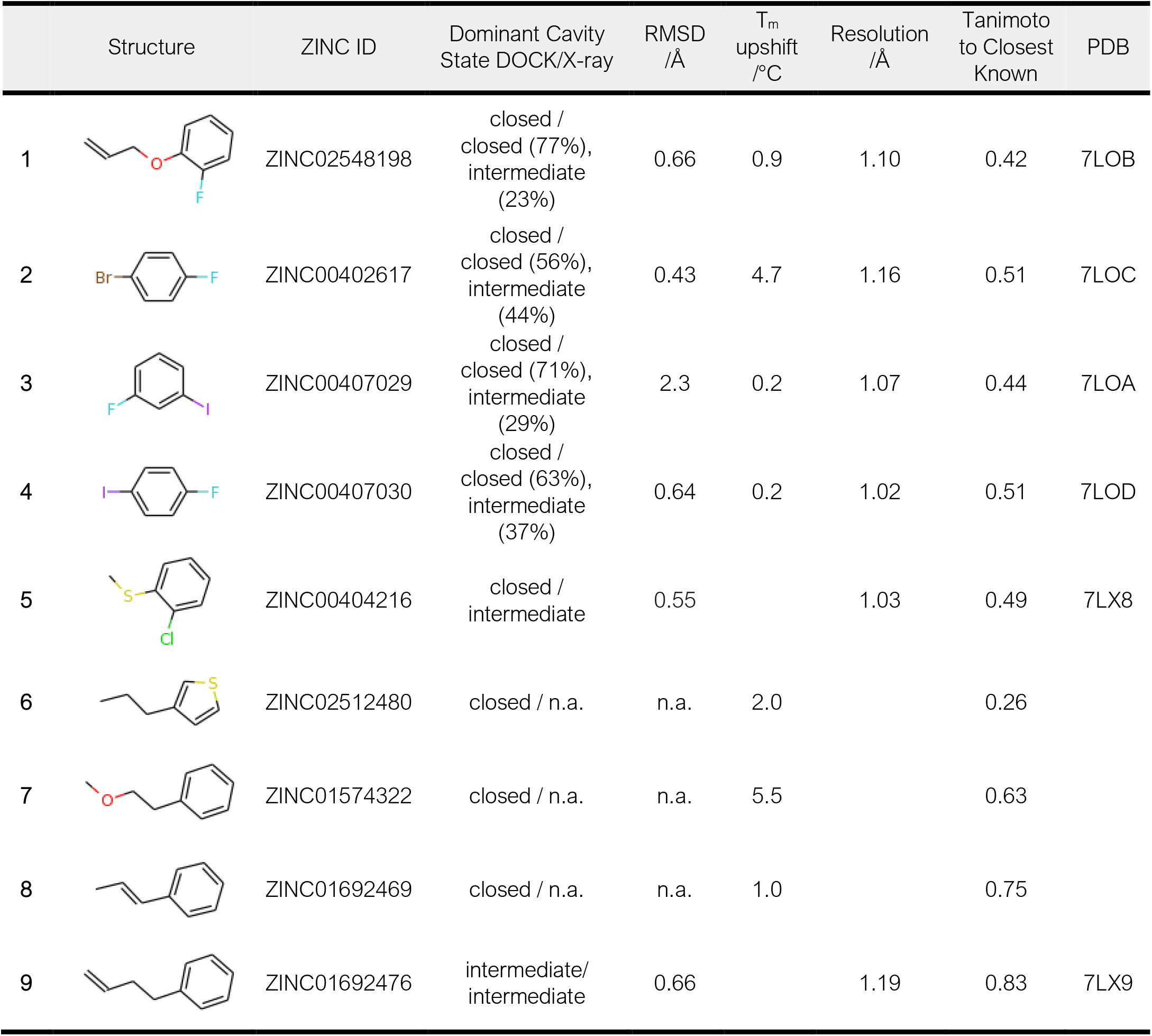
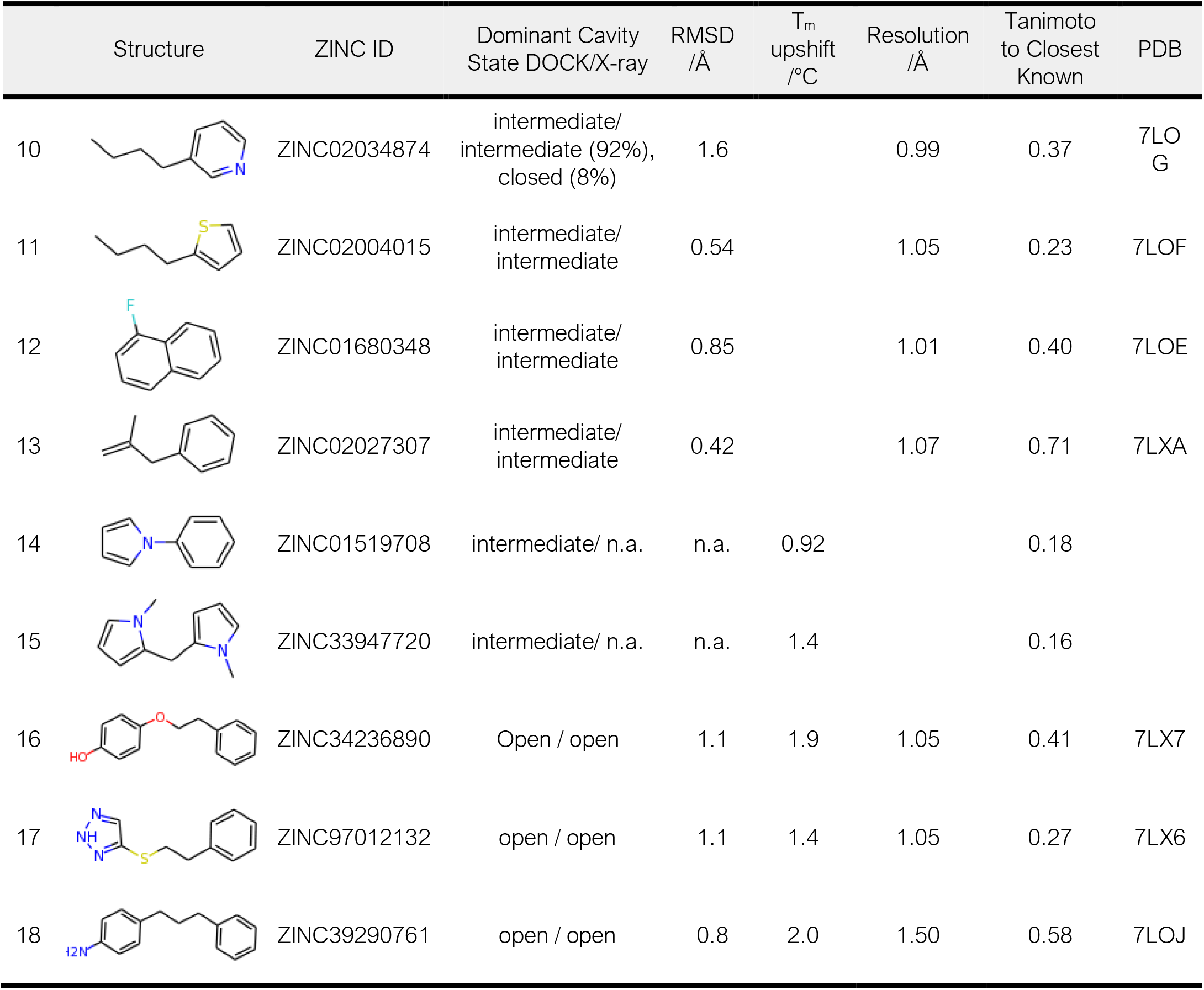
Summary of newly identified L99A binders based on T_m_-upshift and X-ray crystallography

|  | Structure | ZINC ID | Dominant Cavity State DOCK/X-ray | RMSD /Å | T<sub>m</sub> upshift /°C | Resolution /Å | Tanimoto to Closest Known | PDB |
| --- | --- | --- | --- | --- | --- | --- | --- | --- |
| 1 |  | ZINC02548198 | closed / closed (77%), intermediate (23%) | 0.66 | 0.9 | 1.10 | 0.42 | 7LOB |
| 2 |  | ZINC00402617 | closed / closed (56%), intermediate (44%) | 0.43 | 4.7 | 1.16 | 0.51 | 7LOC |
| 3 |  | ZINC00407029 | closed / closed (71%), intermediate (29%) | 2.3 | 0.2 | 1.07 | 0.44 | 7LOA |
| 4 |  | ZINC00407030 | closed / closed (63%), intermediate (37%) | 0.64 | 0.2 | 1.02 | 0.51 | 7LOD |
| 5 |  | ZINC00404216 | closed / intermediate | 0.55 |  | 1.03 | 0.49 | 7LX8 |
| 6 |  | ZINC02512480 | closed / n.a. | n.a. | 2.0 |  | 0.26 |  |
| 7 |  | ZINC01574322 | closed / n.a. | n.a. | 5.5 |  | 0.63 |  |
| 8 |  | ZINC01692469 | closed / n.a. | n.a. | 1.0 |  | 0.75 |  |
| 9 |  | ZINC01692476 | intermediate/intermediate | 0.66 |  | 1.19 | 0.83 | 7LX9 |

**Table 1.** continued.
|  | Structure | ZINC ID | Dominant Cavity State DOCK/X-ray | RMSD /Å | T<sub>m</sub> upshift /°C | Resolution /Å | Tanimoto to Closest Known | PDB |
| --- | --- | --- | --- | --- | --- | --- | --- | --- |
| 10 |  | ZINC02034874 | intermediate/<br>intermediate (92%),<br>closed (8%) | 1.6 |  | 0.99 | 0.37 | 7LOG |
| 11 |  | ZINC02004015 | intermediate/<br>intermediate | 0.54 |  | 1.05 | 0.23 | 7LOF |
| 12 |  | ZINC01680348 | intermediate/<br>intermediate | 0.85 |  | 1.01 | 0.40 | 7LOE |
| 13 |  | ZINC02027307 | intermediate/<br>intermediate | 0.42 |  | 1.07 | 0.71 | 7LXA |
| 14 |  | ZINC01519708 | intermediate/ n.a. | n.a. | 0.92 |  | 0.18 |  |
| 15 |  | ZINC33947720 | intermediate/ n.a. | n.a. | 1.4 |  | 0.16 |  |
| 16 |  | ZINC34236890 | Open / open | 1.1 | 1.9 | 1.05 | 0.41 | 7LX7 |
| 17 |  | ZINC97012132 | open / open | 1.1 | 1.4 | 1.05 | 0.27 | 7LX6 |
| 18 |  | ZINC39290761 | open / open | 0.8 | 2.0 | 1.50 | 0.58 | 7LOJ |

For 13 of these ligands we determined X-ray crystal structures bound to L99A (**Table 1, SI Figure S6, SI Table 3**). Crystals diffracted from 0.99 to 1.5 Å resolution. In all 13 of the crystal structures, we can see the unrefined difference electron density maps (F_o_-F_c_) clearly defining both ligand positions and that of the F-helix, whose opening defines the three conformational states of the cavity. Crystals diffracted from 0.99 to 1.5 Å resolution. In all 13 of the crystal structures, we can see the unrefined difference electron density maps (Fo-Fc) clearly defining both ligand positions and that of the F-helix, whose opening defines the three conformational states of the cavity (**SI Figure S6**). The quality of these observations made us confident on our ability to predict these structures. Of the five closed state structures, four (PDB ID’s 7LOB, 7LOC, 7LOA and 7LOD) also had the alternate electron density for the intermediate state in the F-helix, thus both intermediate and closed conformation existed in the same structures with the dominant conformation being closed. For PDB 7LOB, the closed cavity conformation occupied 77% of the observed electron density while the rest was modeled as intermediate. The percentage of closed cavity conformation for the other three closed state structures with PDB ID’s 7LOC, 7LOA and 7LOD were 56%, 71% and 63% respectively. The only intermediate structure that also had an alternate conformation for closed state included PDB ID 7LOG with 92% of the conformation being intermediate and the rest modeled as closed state. (Alternate conformational states with less than 8% occupancy were not modeled in the crystal structures.) Encouragingly, the docked poses of the predicted ligands closely resembled the crystallographic geometries, with RMS deviations ranging from 0.42 to 2.3 Å, and a mean RMSD of 0.90 Å (**Figure 4**). Crucially, all ligands but one (compound **5**) bound to the conformational state for which it had been predicted in the weighted-ensemble docking (**Table 1**). Thus, ligands predicted to bind to the closed state, compounds **1** through **4**, were observed to bind to that state, the seven ligands predicted to bind to the intermediate state, compounds **9** through **13**, bound to it, and the three ligands predicted to bind to the open state, compounds **16**, **17** and **19** bound to it. Only compound **5**, which was predicted to bind to the closed state, was found to bind to the intermediate state. For all of these complexes, there was little ambiguity in making these state assignments—in the four closed state structures, the F-helix adopted a classic α-helical geometry, with residues V111 and A112 adopting a “down” conformation, into the site, and no pathway between the cavity and bulk solvent—the ligands were completely enclosed by the residues defining the cavity. For the six intermediate state structures, residues T109 to G113 shift “upwards”, with Val111 rotating about its *χ*_1_ angle, enlarging the cavity. Finally, in all three open state structures, the F-helix adopts a 3-10 helix, and a clear channel opens up to solvent, which the docked ligands exploit. Whereas the docking hit-rate for this state was relatively low, all three ligands bound in geometries that corresponded closely to the docking predictions (**Figure 4L, M, and N**). Clearly, the driving force for binding in the closed, intermediate, and open states was sterics, but in the open state there is also a role for the orientation of polar groups, which typically interacted with the backbone of residue G107 in the unwound F-helix and also with bulk solvent. Encouragingly, these interactions were captured in the docking.

## Discussion

Three key results emerge from this study: **first,** MD simulations, including accelerated MD (aMD), can access and energy-weight alternative conformational states of binding sites; **second,** these weighted conformations substantially improve docking hit rates versus unweighted states; and **third** the docking hits predicted for each state actually bind to those states. Using MD—and, indeed, other methods, such as simply relying on experimental observables—to access and weight protein conformations for docking has confronted two problems—accessing the correct states, and weighting them by energy. In previous work, even when states are sampled correct, docking can be biased by those states that best fit ligands, which are often those that are higher in energy. These high energy states thus act as decoys, crowding out more favorable solutions. Drawing on experimental observables, several studies have shown that when states are properly weighted by conformational energies, docking results improve (4, 9). However, depending only on experimental observables, such as multiple state occupancies, restricts this approach to the relatively few targets that afford such high-quality observables. Thus, a crucial result of this study is that both conventional MD (cMD) and, encouragingly, also aMD can both sample experimentally accessible states and usefully weight them (**Figure 2A**). Tested retrospectively in the model cavity site L99A, docking hits are dominated by the highest energy open conformation of the site, which is the one that can accommodate the largest ligands. With the aMD-derived population weights applied, retrospective hit rates improve substantially (**Figure 2C-E**). More compellingly, in prospective screens, a high 55% hit rate is found for new ligands, topologically different from those previously known (**Table 1**). The ability to readily determine structures with L99A allowed us to interrogate these new ligands at atomic resolution. For 12 out of 13 structures determined, each new ligand predicted bound to the state for which it was predicted (**Figure 4**).

Besides its high sampling efficiency compared to cMD, an advantage of the aMD approach is its pathway independence (62). Hence, no prior knowledge of a systems free energy landscape needs to be incorporated in the form of a reaction coordinate or collective variable. The required parameters can be derived from energetic averages of short cMD simulations. The workflow we present can thus be applied to any system of interest. Within this work we focus on benchmarking the reliability of aMD-derived energy weights, which we then apply on experimental holo structures. However, it has been shown previously that MD simulations are also apt to suggest protein conformations for ligand discovery, which have not been observed before (8, 61). Selecting single structures from thousands of snapshots visited within MD trajectories is a tedious and typically ambiguous challenge. The kinetic clustering and coarse-graining embedded in a typical MSM workflow represents a most robust and straight-forward approach to this problem. The combination of enhanced and conventional MD simulations (**Figure 1A-B**) may be an efficient method for extensive phase space exploration, with ultimately unbiased transition rates, and thus may offer reliable energetic weightings for the sampled states. This workflow should be transferrable to most biomolecular system.

The energy weights, which we derived from the aMD simulations, substantially improved our docking hit rates, both for retrospective and prospective screens. We find that the energetic penalty term is crucial, in particular for the high-energy open state. Without accounting for the energetic cost of the cavity opening, large decoy molecules dominate the top ranks of the retrospective docking (**Figure 2D**). Larger ligands are inherently favored by the docking scoring function, for this and for other sites, simply because they can make more interactions. Incorporating energy weights helps balance the ligands and states among the top-ranked molecules. While similar observations have been made based on high quality experimental weighting, an advantage of the MD approaches investigated here is that they should be applicable to the majority of targets for which such weighted states are not available directly from experimental observables.

Encouragingly, predicted and experimental binding site conformations corresponded well. For 12 of 13 ligand-bound structures, the flexible receptor docking predicted the correct dominant receptor conformation. This is most compelling for the open-state, where the site doubles in volume and opens it to the bulk solvent, making it substantially more complicated in terms of its potential interactions and biophysics. The three molecules that bind to this conformation, compounds **16, 17**, and **18**, include polar groups such as a phenolic hydroxyl, a triazole, and an amine (**Figure 4 L-N**), which hydrogen-bond with the backbone of G107 in the F-helix, a group previously unavailable to them in the closed and intermediate conformations. In the open state, these groups are also open to bulk solvent, which likely has a role in reducing the desolvation penalty they would otherwise pay. Balanced against the ability to predict this state for these ligands, and to predict the ligand-bound geometries with high fidelity, was the relatively low hit rate for molecules predicted to bind to the open state. This at least partly reflects the loss of the steric constraint that effectively separates ligands from many larger non-binders in the much smaller closed and intermediate cavity conformations—in the open state, the L99A “cavity” begins to reflect the more open sites typical of drug targets, as do its docking hit rates (3/11). Likely also contributing was the relatively low solubility of the larger molecules versus smaller ligands for a site whose maximal affinity is close to 10^-4^ M.

Certain limitations of this study merit airing. Whereas we believe that aMD can be used to not only weight but sample relevant conformations, here we relied on crystallographically defined conformational states, from which we calculated MD-based probabilities. Until this is demonstrated prospectively, a substantial undertaking, the generalizability of this approach will remain tentative. We do note that MSM-based state definitions have shown much promise in ligand discovery (8). Naturally, this study was conducted in a model system, the apolar cavity site of L99A, that intentionally simplifies the problem, eliminating many challenges typically faced in docking (e.g., a usually heterogenous recognition surface, partly exposed to solvent, is radically simplified). The hit rates experienced here, and the sampling challenges, will be increased in more complicated, drug-relevant binding sites. By the same standard, the simplifications afforded by L99A, like all model systems, allowed us to focus on the challenge at hand, sampling and including multiple protein conformations. Finally, we note that though the overall hit rate for the docking was high, at 55%, the hit rate for the more challenging open cavity, though not disreputable at 27%, was substantially lower, reflecting the challenges emerging as one moves from an enclosed cavity to the more typical binding sites. The lower hit rate reflects well-known problems with docking, we believe, not really challenges attending the weighted ensemble approach; all but one of the ligands predicted, after all, bound in the states to which they were docked.

These caveats should not distract from the central observations of this study – molecular dynamics simulations, including the resource-efficient accelerated MD, can sample relevant states and usefully energy-weight them. The resulting energy penalties, incorporated into multi-conformer receptor docking, allow one to access a much broader range of chemotypes without being dominated by high-energy structures. The approach has the promise of generality, and potentially may be applied to the vast number of systems where such states are unavailable from experimental data, but are likely to play a key role in the success of large library docking screens.

## Materials and Methods

### MD Simulation Setup and Analysis

Structures, coordinates and topologies for the MD simulations were prepared using modules implemented in AMBER14 (63). The analysis was performed using in-house python scripts, cpptraj (64) and PyEMMA 2.5.7 (65). Further details are available in the Supplementary Information.

### Flexible Receptor Docking

All library screens were performed using the flexible receptor docking protocol, scripts, and programs implemented in DOCK3.7 (66). A detailed description is included in the Supplementary Information

### Protein Purification

T4 Lysozyme L99A was cloned as previously described (67). Briefly, pET-29 plasmid (EMD Biosciences, Darmstadt) containing the gene L99A T4 lysozyme gene was transformed into *E. coli* BL21(DE3) cells. The C-terminally hexa-histidine tagged L99A T4 lysozyme was expressed in BL21 cells for 4 hours following induction with 0.5 mM IPTG overnight at 18° C in presence of kanamycin. Cells were harvested by centrifugation at 4000 rpm for 20 minutes followed by resuspension of cells in lysis buffer containing 20 mM HEPES pH 6.8, 10 mM Imidazole, 5 mM beta-mercaptoethanol and lysed using a sonicator. Following lysis, the His-tagged protein was bound to Ni-NTA agarose and subjected to wash with buffer containing 20 mM HEPES pH 6.8, 20 mM Imidazole and 5 mM beta-mercaptoethanol. Finally, the protein was eluted using elute buffer containing 20 mM HEPES pH 6.8, 250 mM imidazole and 5 mM β-mercaptoethanol. The eluted protein was dialyzed in 200 mM KCl, 5 mM β-mercaptoethanol and 50 mM phosphate buffer pH 6.6 and concentrated to 20 mg/ml before flash freezing in liquid nitrogen to store the protein at −80° C.

### Differential Scanning Fluorimetry (DSF)

T4 Lysozyme L99A was incubated with SYPRO Orange dye (Thermo Fisher, S6650) in a 384-well PCR microplate (VWR, 10011-194), with a final volume of 15 μL per well, including 2.5 μM T4Lys, and 2.5× Sypro dye in 50 mM KPi, pH 6.5. The temperature was ramped from 30 to 95 °C at a rate of 1 °C/min and fluorescence of the dye monitored by qPCR machine. Melting temperatures were determined by the Life Cycler Thermal Shift Analysis software. L99A was mixed with a final 100 micromolar concentration of small molecules prior to DSF. The difference in the melting temperature was calculated for each small molecule-L99A mixture.

### Protein Crystallization

Crystals for T4 lysozyme L99A were set at a protein concentration of 10 mg/ml using vapor diffusion hanging drop method. Crystals grew overnight at 20° C in buffer containing 0.1 M Tris pH 8.0, 22% PEG, 4% Isopropanol, 50 mM BME and 50 mM 2-hydroxyethyl disulfide. Ligands were soaked in the crystals and crystals were left overnight at 20° C before cryocooling them in 25% ethylene glycol for data collection.

### Structure determination and Refinement

L99A-ligand datasets were collected at beamline 8.3.1 of the Advanced Light Source (ALS, Lawrence Berkeley Lab, CA) with wavelength of 0.95386 and a temperature of 100K. All datasets belonged to P3_2_2_2_1 with one molecule in the asymmetric unit. The datasets were processed, scaled, and merged using XDS and AIMLESS. MOLREP was used for molecular replacement using the protein model from PDB ID 4W57. The F-helix residues 107-115 and ligand were removed from PDB ID 4W57 during molecular replacement giving unbiased electron density for the ligands and F-helix in the initial electron density maps. Geometry restraints of ligands were created in eLBOW-PHENIX. Initial model fitting and addition of waters was done in COOT [109] followed by refinement in REFMAC [99]. Following modeling of the ligand in COOT, several rounds of refinement were carried out using PHENIX. For each structure, geometry was assessed using Molprobity and PHENIX polygon. Structures were not compared until completion of the refinement for each ligand to prevent biasing of refinement of different states of lysozyme. Datasets have been deposited to the PDB as 7LOA, 7LOB, 7LOC, 7LOD, 7LOE, 7LOF, 7LOG, 7LOJ, 7LXA, 7LX6. 7LX7, 7LX8, 7LX9.

## Acknowledgements

We thank Taia Wu and Jason Gestwiki for assistance with the DSF fluorimetry. We thank Brian Bender, Matt Smith, and Ying Yang for reading this manuscript. Furthermore, we thank Marcus Fischer for helpful discussions. Supported by R35GM122481 (to BKS), by the Austrian Science Fund (FWF) via grant P30737 (to KRL) and by a Marshall Plan and a Marietta Blau fellowship (to ASK).

## Supplementary Information

### Supplementary Materials and Methods

#### Molecular Dynamics Simulation Setup and Analysis

For exhaustive exploration of L99A’s conformational space accelerated molecular dynamics (aMD) simulations were performed. The simulations were started from an open state structure (PDB: 4W59), where we removed the ligand n-hexylbenzene from the cavity, as well as all crystallization agent and water molecules. With the LEaP module implemented in AMBER14 (1), we created topologies and initial coordinate files using the AMBER 99SB-ILDN force field (2). The protein was solvated with a truncated octahedral box of TIP3P water molecules(3) and a minimum wall distance of 12 Å. We further apply an exhaustive equilibration protocol to relax the system in an NPT ensemble prior to productive simulation runs (4). The aMD specific parameters, i.e., threshold energy and boosting parameter, were determined from the final short cMD simulation as described previously (**SI Table S1**) (5, 6).

To maximize computational efficiency all simulations were performed with the GPU implementation of AMBER14s pmemd module (7). Hence, the particle-mesh Ewald (PME) method was used to treat long ranging electrostatic interactions and a non-bonded cutoff of 8 Å (8). We apply a Langevin thermostat (9) with a collision frequency of 2 ps-1 to maintain a simulation temperature of 300K and a Berendsen barostat (10) with a relaxation time of 2 ps to simulate constant atmospheric pressure. To allow for a the timestep of 2 fs all bonds involving hydrogen atoms were constraint using the SHAKE algorithm (11).

We performed five replica aMD simulations of 100 ns length starting from the same coordinates with different velocities. The accumulated simulation time of 500 ns was then clustered using the hierarchical agglomerative clustering implemented in cpptraj (12) using average linkage and a cutoff distance of 0.8 Å. We assigned the resulting clusters to the open, intermediate or closed cavity state based on structural similarity of the representative structure. In **SI Figure S1** we visualize the opening and closing of the cavity color-coded according to the state-populations derived from the clustering of the combined trajectory. The cluster populations where then reweighted with an approximation of the exponential term using a Maclaurin series of the 10th order (13). To estimate the uncertainty of these reweighted state populations we applied the same procedure for each 100 ns aMD trajectory individually (**SI Table S2**).

Furthermore, we performed extensive sampling with cMD simulations to derive unbiased thermodynamic and kinetic information with the aid of an MSM. On the one hand, we seed cMD simulations from the aMD ensemble. On the other hand, cMD simulations were started from an open state crystal structure, as described above, to increase the statistical robustness. In total we accumulated 7.75 μs of simulation time.

From the unbiased cMD simulation data we constructed an MSM using PyEMMA 2.5.7.(14) Based on structural characteristics in the ligand-bound crystal structures, we selected the distance between the buried residue ALA99 and the Cα atoms of the F-helix as well as the rotation of residue V112 (chi1) as input features a time-lagged independent component analysis (TICA)(15) with a lag-time of 0.5 ns. After projecting the structural information, we divide the TICA space into 100 microstates using a k-means clustering. Based on the discretized trajectory we then built a Bayesian MSM (16) with a lag time of 0.5 ns. We perform a Perron-cluster cluster analysis+ (PCCA+) to coarse-grain our model into four states, as deduced from gap between successive eigenvalues (**SI Figure S2**). A Chapman-Kolmogorov test displaying the robustness of the model is shown in **SI Figure S3**. A visualization of the thermodynamics and kinetics calculated from the MSM is depicted in the **SI Figure S4**.

#### Flexible Receptor Docking

The flexible receptor docking protocol, scripts, and programs implemented in DOCK3.7 were used to calculate and score ligand poses with each receptor conformation (17). The crystal structure of n-butyl-benzene bound to L99A (PDB 4W57) was prepared using REDUCE (18) to add hydrogens. For the orientation of docked ligands in the binding site, the crystallographic ligand atoms were converted into “spheres”, which are pseuodoatoms that are used to orient new docked ligands (19). The generation of these spheres was accomplished as implemented in the Blastermaster script distributed with DOCK3.7 using the SPHGEN program. Using QNIFFT (20), electrostatic potentials were calculated by solving the Poisson-Boltzmann equation, and stored on a lattice for scoring look-up. Van der Waals interactions were calculated using CHEMGRID (21), and ligand desolvation grids were calculated using SOLVMAP (22). Retrospective testing was based on 68 known ligands (23–29), for which we generated property-matched decoys using the DUD-E (30) workflow. For prospective screening, we docked a library of 985,201 molecules from a subset of ZINC15 (31) with a cLogP up to 4 and a molecular weight up to 300 Da.

### Supplementary Figures and Tables

**Supplementary Table S 1.** Parameters applied in aMD simulations as calculated from average energies, particles and residue number.

|  |  |
| --- | --- |
| $E_{\text{dihed}}$ | 2382 |
| $\alpha_{\text{dihed}}$ | 66 |
| $E_{\text{tot}}$ | -80587 |
| $\alpha_{\text{tot}}$ | 4189 |

**Supplementary Figure S 1.**
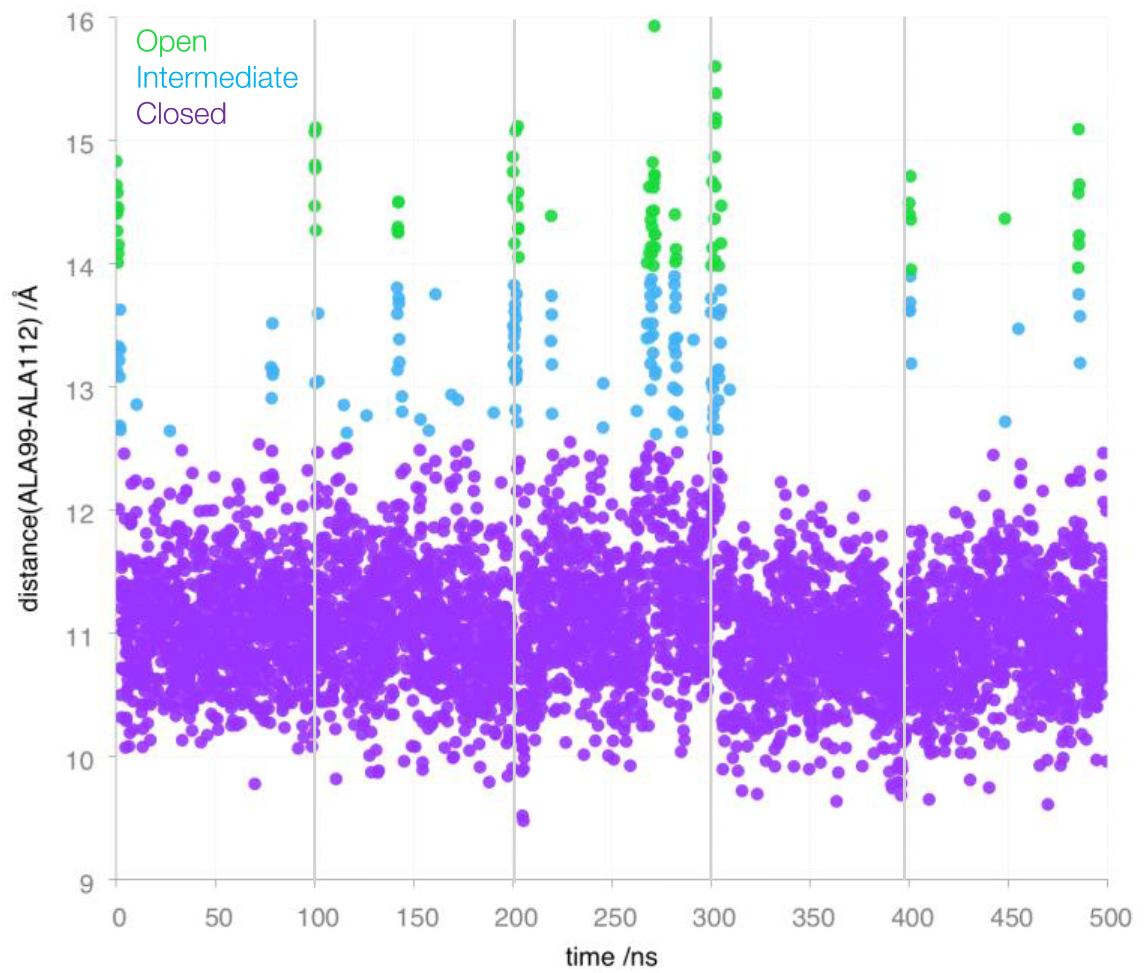
Cavity opening and closing during aMD simulations. The distance between the bottom of the cavity (ALA99) and ALA112 in the F-helix is colored according to the conformational state as defined by the clustering.

**Supplementary Table S 2.** Error estimation of state populations. The reweighted population of closed, intermediate and open state was calculated from five 100 ns aMD trajectories individually to estimate the associated uncertainty.

| State | aMD1 | aMD2 | aMD3 | aMD4 | aMD5 | Average | Standard Deviation |
| --- | --- | --- | --- | --- | --- | --- | --- |
| Closed | 96.7% | 97.6% | 95.4% | 97.3% | 98.7% | 97.1% | $\pm 1.2\%$ |
| Intermediate | 2.8% | 1.8% | 3.8% | 1.4% | 1.2% | 2.2% | $\pm 1.1\%$ |
| Open | 0.5% | 0.6% | 0.9% | 1.2% | 0.1% | 0.7% | $\pm 0.4\%$ |

**Supplementary Figure S 2.**
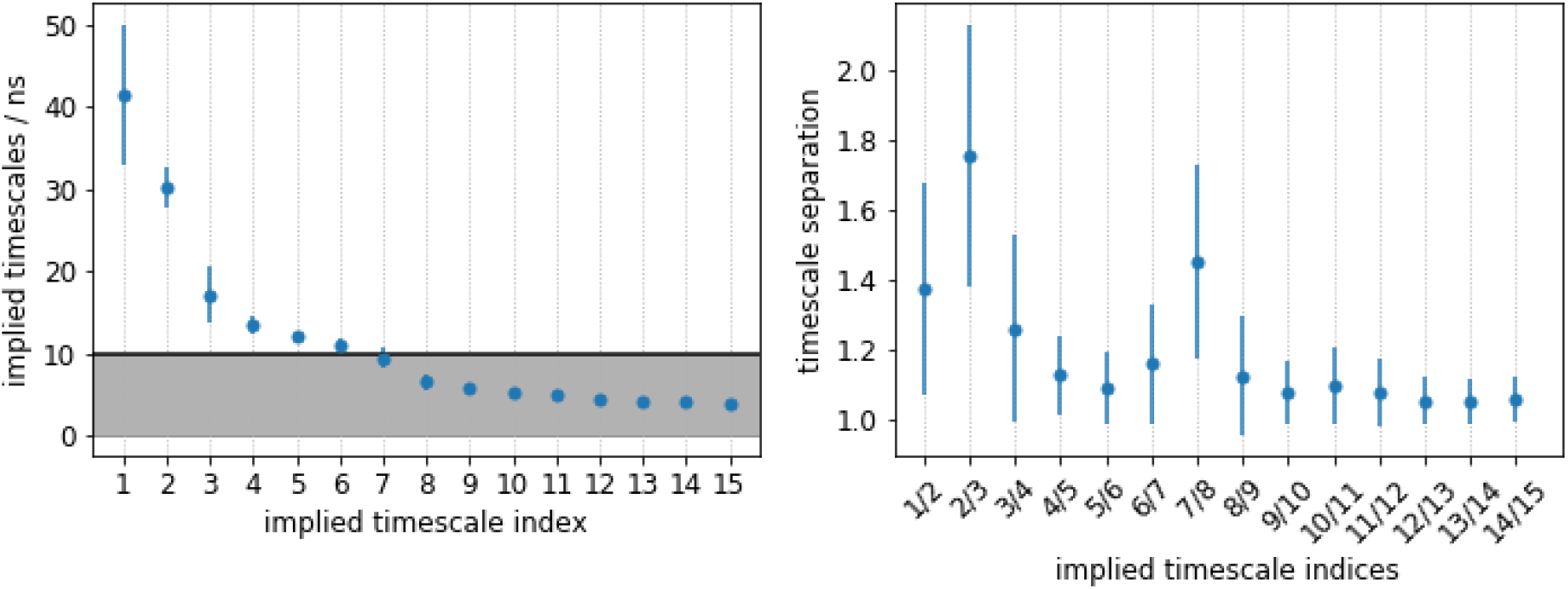
Successive Eigenvalues of the TICA suggesting four macrostates for the PCCA+ coarse-graining of the MSM.

**Supplementary Figure S 3.**
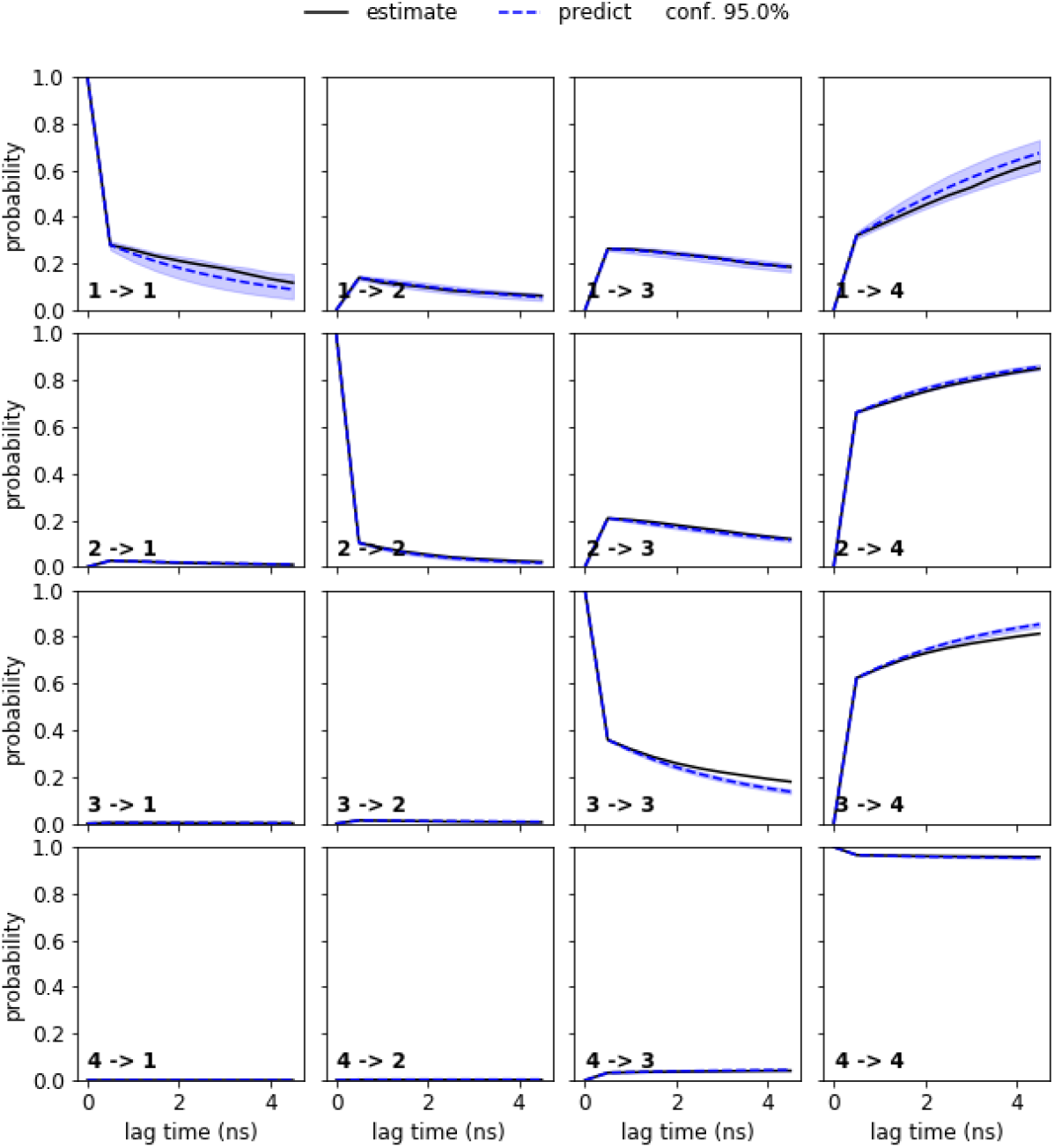
Chapman-Kolmogorov test of the MSM. The test is depicting good agreement between predicted results using the applied lag time of 0.5 ns (dashed line) and estimated at lag times up to 5ns (continuous line).

**Supplementary Figure S 4.**
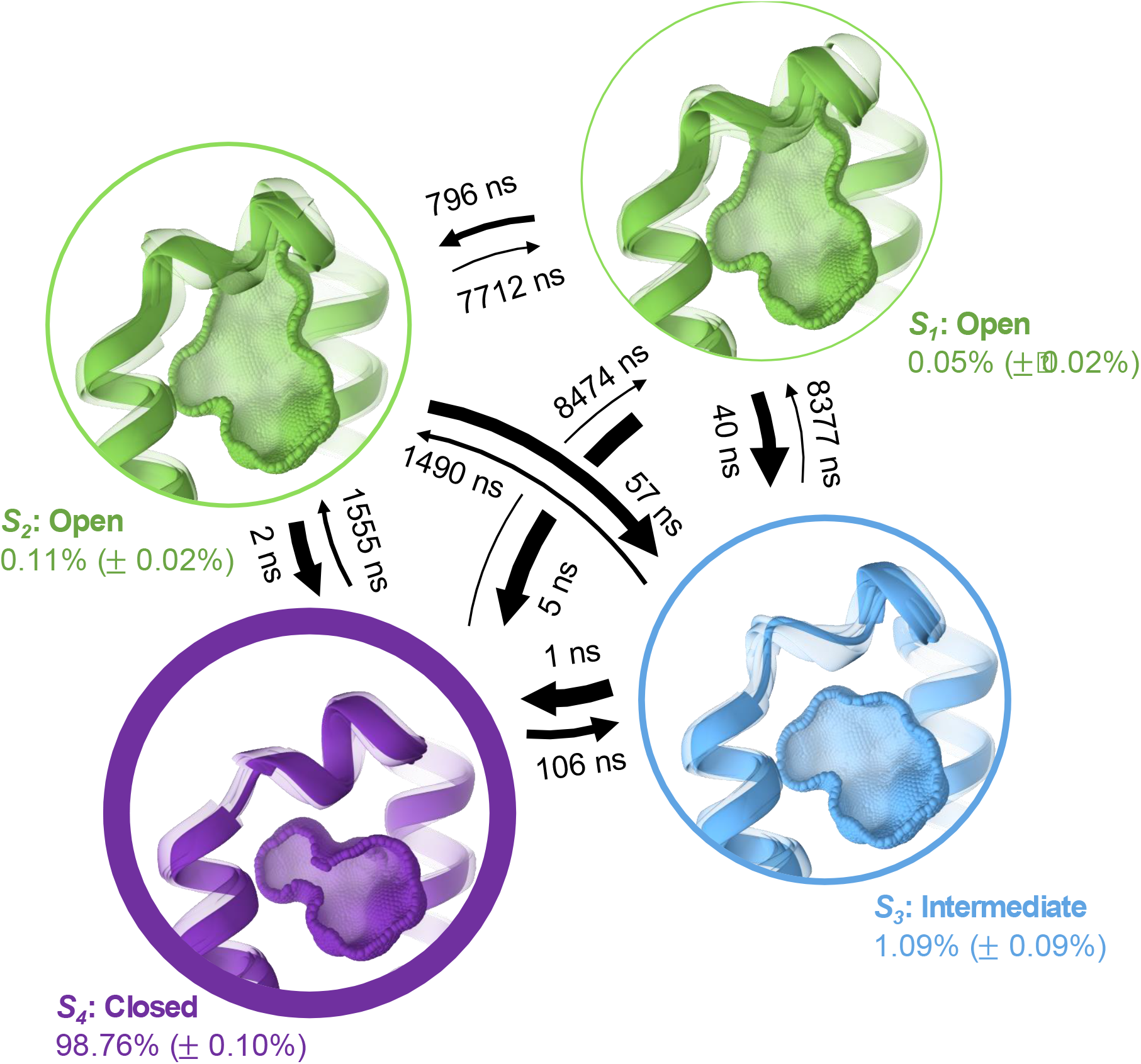
Distinct conformational states of the ligand binding site in L99A. A Bayesian Markov state model of the apo protein ensemble estimates the population of each state in the absence of a ligand. The MSM states S1 (0.05 %) and S2 (0.11 %) both resemble the open state, S3 (1.09 %) represents an intermediate conformational state and the closed conformation is characterized by S4 (98.76 %).

**Supplementary Figure S 5.**
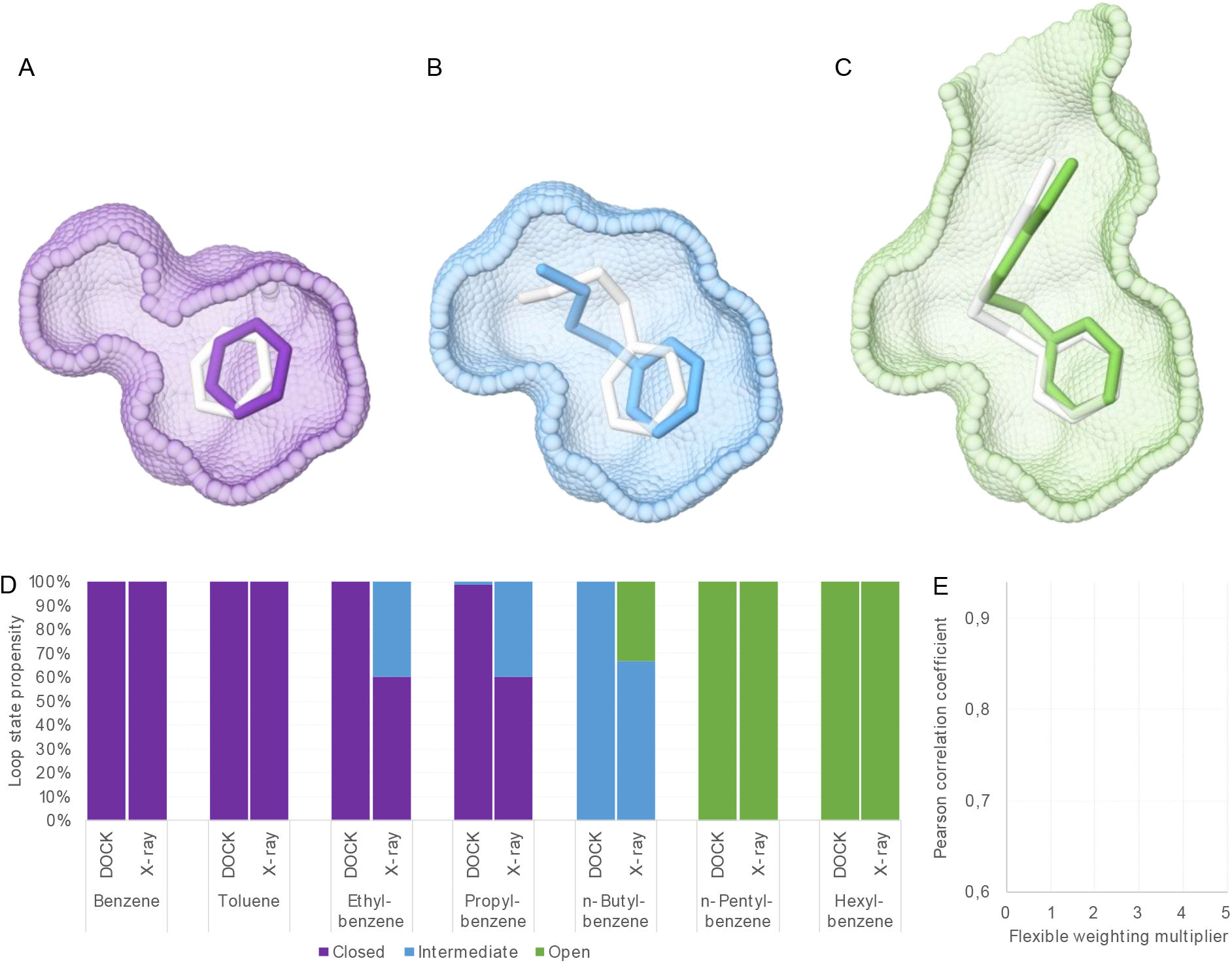
Crystallographic and predicted geometries of known ligands and loop state probabilities. Docking poses for (A) Benzene, a closed state binder (purple), (B) n-Butylbenzene, an intermediate state binder (blue), and (C) n-Hexylbenzene, an open state binder (green) in overlay with the crystallographic ligand geometries (white). (D) State probabilities predicted from DOCK compared to crystallographic occupancies for a homologous ligand series. (E) Pearson correlation coefficient of crystallographic occupancies and state probabilities calculated from the DOCK score as function of the flexible weighting multiplier.

**Supplementary Figure S 6.**
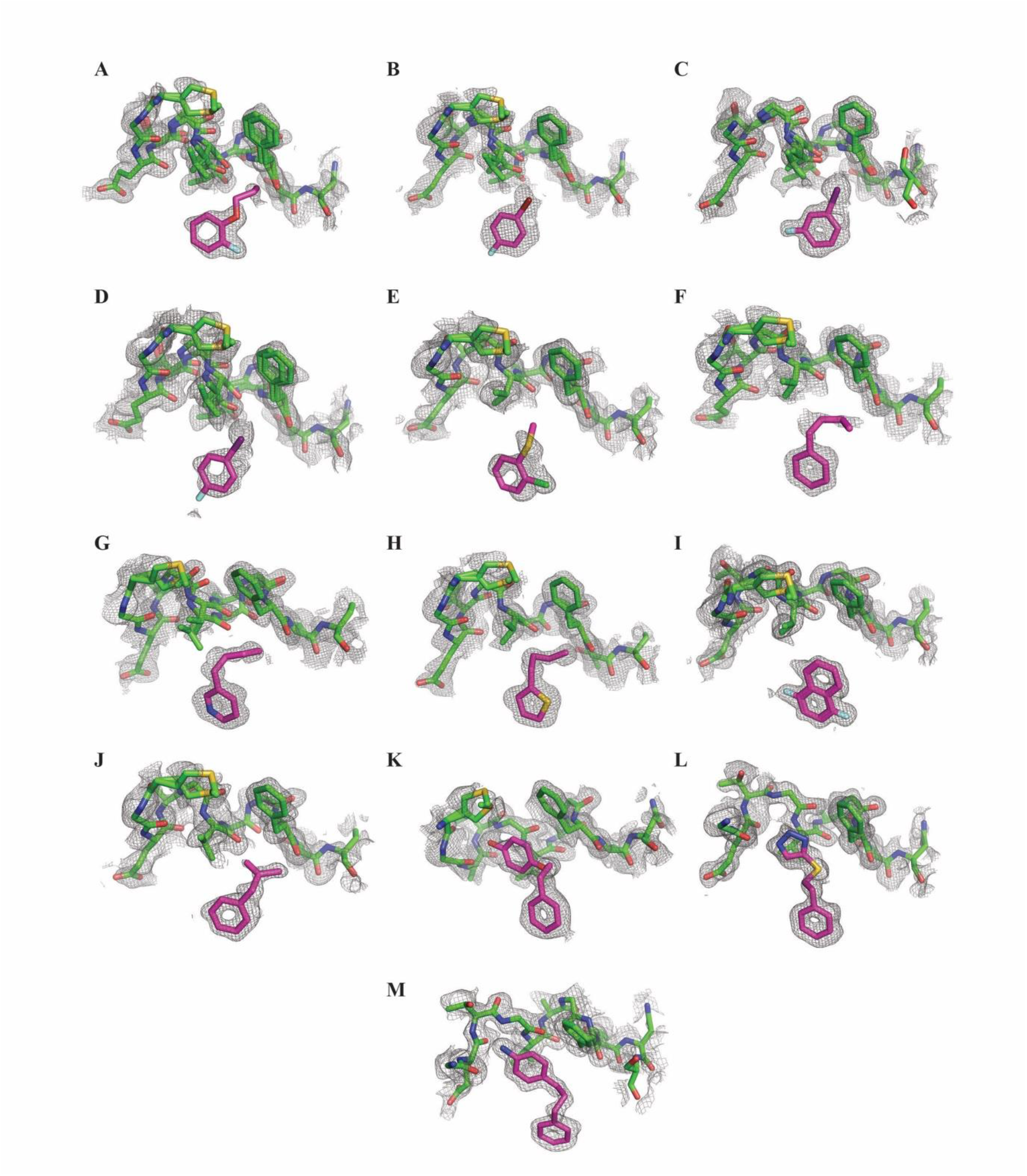
Electron Density Maps for L99A binders and F-helix. The initial Fo-Fc electron density map contoured at 1.6σ around the ligand and F-helix (density in grey) for L99A lysozyme-ligand complexes A. 7LOB, B. 7LOC, C. 7LOA, D. 7LOD, E. 7LX8, F. 7LX9, G. 7LOG, H. 7LOF, I. 7LOE, J. 7LXA, K. 7LX7, L. 7LX6 and M. 7LOJ. Ligand carbons in magenta and protein carbons in green, oxygens red, nitrogens blue, sulfurs yellow and chlorides green.

**Supplementary Table S 3.** Crystallographic Statistics

| **PDB ID** | **7LOB** | **7LOC** | **7LOA** |
| --- | --- | --- | --- |
| **Data collection** |  |  |  |
| Space group | P 32 2 1 | P 32 2 1 | P 32 2 1 |
| **Cell dimensions** |  |  |  |
| *a*, *b*, *c* (Å) | 60.412, 60.412, 96.278 | 60.266, 60.266, 96.411 | 60.057, 60.057, 96.276 |
| *a*, *b*, *g* (°) | 90.00, 90.00, 120.00 | 90.00, 90.00, 120.00 | 90.00, 90.00, 120.00 |
| Resolution (Å) | 45.97 - 1.1 (1.139 - 1.1)* | 52.19 - 1.16 (1.202 - 1.16) | 52.01 - 1.07 (1.108 - 1.07) |
| Total reflections | 1607952 (151824) | 140805 (13413) | 177215 (17042) |
| Unique reflections | 83069 (8240) | 70437 (6741) | 88627 (8536) |
| Multiplicity | 19.4 (18.4) | 2.0 (2.0) | 2.0 (2.0) |
| *R*<sub>sym</sub> or *R*<sub>merge</sub> | 0.08578 (4.381) | 0.01331 (0.4135) | 0.01322 (0.6274) |
| *I* / *sI* | 20.07 (0.69) | 25.08 (1.88) | 24.17 (1.24) |
| Completeness (%) | 99.89 (99.41) | 99.69 (96.98) | 99.43 (96.73) |
| CC1/2 | 1 (0.391) | 1 (0.733) | 1 (0.527) |
| **Refinement** |  |  |  |
| Resolution (Å) | 52.318 - 1.100 | 52.192 - 1.160 | 52.011 - 1.070 |
| Reflections used in refinement | 82977 (8192) | 70424 (6737) | 88609 (8529) |
| Reflections used for R-free | 4176 (449) | 3528 (338) | 4481 (435) |
| R-work/R-free (%) | 20.90/22.18 | 19.76/20.80 | 20.76/21.25 |
| **No. atoms** |  |  |  |
| Protein | 1332 | 1345 | 1353 |
| Ligand/ion | 23 | 24 | 28 |
| Water | 116 | 124 | 122 |
| **B-factors** |  |  |  |
| Protein | 16,05 | 15,39 | 16,05 |
| Ligand/ion | 21,1 | 24,03 | 24,88 |
| Water | 23,47 | 22,91 | 23,96 |
| **R.m.s. deviations** |  |  |  |
| Bond lengths (Å) | 0,004 | 0,004 | 0,004 |
| Bond angles (°) | 0,76 | 0,75 | 0,78 |

**Supplementary Table S 4.**
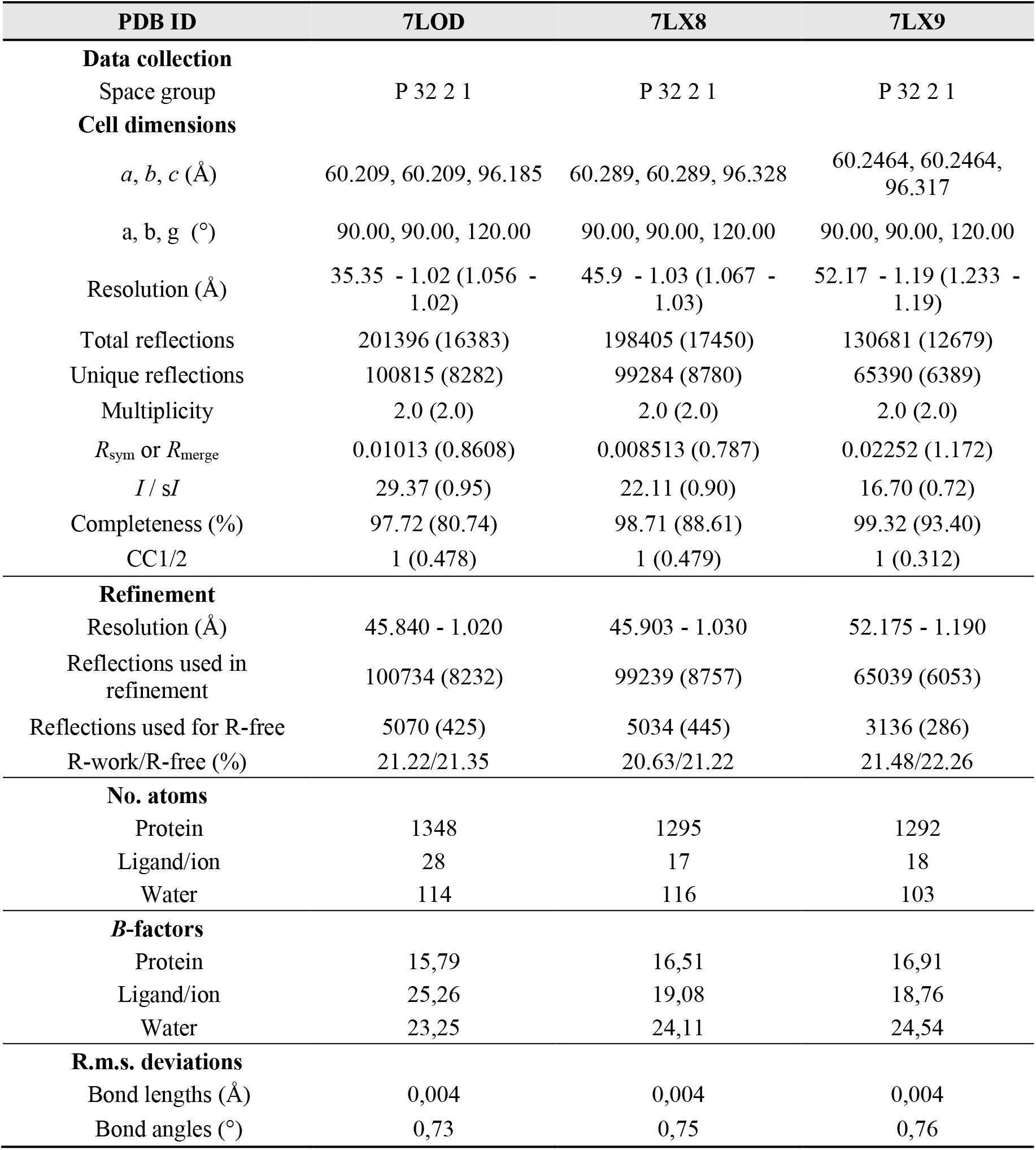

**Supplementary Table S 5.**

| PDB ID | 7LOG | 7LOF | 7LOE |
| --- | --- | --- | --- |
| **Data collection** |  |  |  |
| Space group | P 32 2 1 | P 32 2 1 | P 32 2 1 |
| **Cell dimensions** |  |  |  |
| *a*, *b*, *c* (Å) | 60.2482, 60.2482, 96.119 | 60.181, 60.181, 96.346 | 60.0782, 60.0782, 96.266 |
| *a*, *b*, *g* (°) | 90.00, 90.00, 120.00 | 90.00, 90.00, 120.00 | 90.00, 90.00, 120.00 |
| Resolution (Å) | 45.86 - 0.99 (1.025 - 0.99) | 45.84 - 1.05 (1.088 - 1.05) | 45.77 - 1.01 (1.046 - 1.01) |
| Total reflections | 216941 (16204) | 182471 (14771) | 205681 (16484) |
| Unique reflections | 108752 (8263) | 91335 (7448) | 102989 (8331) |
| Multiplicity | 2.0 (2.0) | 2.0 (2.0) | 2.0 (2.0) |
| *R*<sub>sym</sub> or *R*<sub>merge</sub> | 0.007583 (0.3653) | 0.009087 (0.6067) | 0.00924 (0.7696) |
| *I* / *sI* | 31.03 (2.28) | 20.73 (1.27) | 24.59 (1.08) |
| Completeness (%) | 96.47 (74.36) | 96.43 (79.33) | 97.27 (79.24) |
| CC1/2 | 1 (0.786) | 1 (0.497) | 1 (0.507) |
| **Refinement** |  |  |  |
| Resolution (Å) | 52.176 - 0.990 | 52.118 - 1.050 | 52.029 - 1.010 |
| Reflections used in refinement | 108714 (8253) | 91305 (7421) | 102914 (8286) |
| Reflections used for R-free | 5610 (436) | 4713 (401) | 5258 (439) |
| R-work/R-free (%) | 19.98/20.31 | 19.75/20.06 | 20.39/21.88 |
| **No. atoms** |  |  |  |
| Protein | 1314 | 1304 | 1308 |
| Ligand/ion | 30 | 21 | 34 |
| Water | 151 | 117 | 126 |
| **B-factors** |  |  |  |
| Protein | 13,17 | 15,58 | 13,64 |
| Ligand/ion | 17,32 | 17,48 | 14,84 |
| Water | 20,03 | 22,77 | 21,01 |
| **R.m.s. deviations** |  |  |  |
| Bond lengths (Å) | 0,004 | 0,004 | 0,004 |
| Bond angles (°) | 0,78 | 0,76 | 0,75 |

**Supplementary Table S 6.**
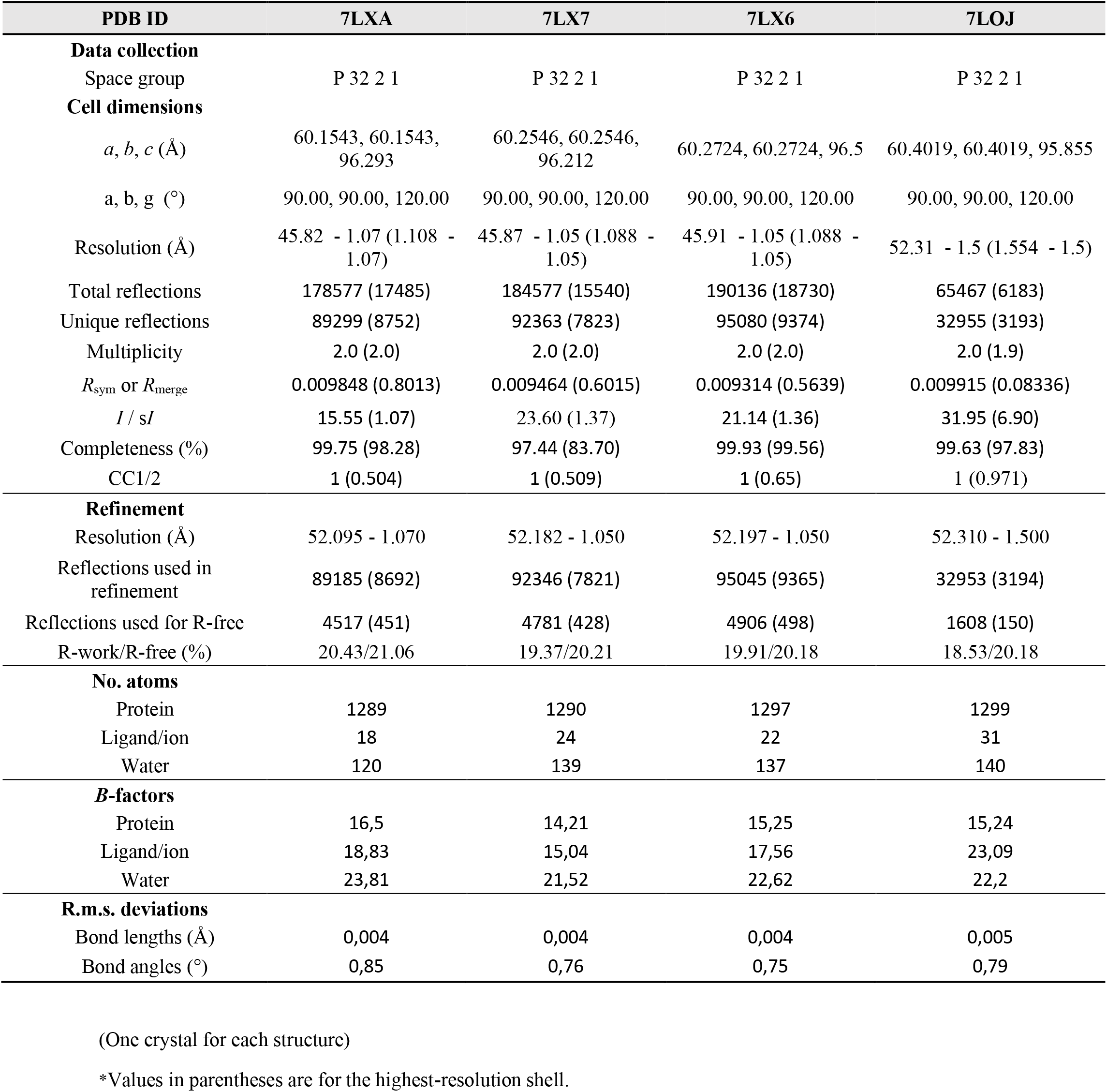

| PDB ID | 7LXA | 7LX7 | 7LX6 | 7LOJ |
| --- | --- | --- | --- | --- |
| **Data collection** |  |  |  |  |
| Space group | P 32 2 1 | P 32 2 1 | P 32 2 1 | P 32 2 1 |
| **Cell dimensions** |  |  |  |  |
| *a*, *b*, *c* (Å) | 60.1543, 60.1543, 96.293 | 60.2546, 60.2546, 96.212 | 60.2724, 60.2724, 96.5 | 60.4019, 60.4019, 95.855 |
| *a*, *b*, *g* (°) | 90.00, 90.00, 120.00 | 90.00, 90.00, 120.00 | 90.00, 90.00, 120.00 | 90.00, 90.00, 120.00 |
| Resolution (Å) | 45.82 - 1.07 (1.108 - 1.07) | 45.87 - 1.05 (1.088 - 1.05) | 45.91 - 1.05 (1.088 - 1.05) | 52.31 - 1.5 (1.554 - 1.5) |
| Total reflections | 178577 (17485) | 184577 (15540) | 190136 (18730) | 65467 (6183) |
| Unique reflections | 89299 (8752) | 92363 (7823) | 95080 (9374) | 32955 (3193) |
| Multiplicity | 2.0 (2.0) | 2.0 (2.0) | 2.0 (2.0) | 2.0 (1.9) |
| *R*<sub>sym</sub> or *R*<sub>merge</sub> | 0.009848 (0.8013) | 0.009464 (0.6015) | 0.009314 (0.5639) | 0.009915 (0.08336) |
| *I* / *sI* | 15.55 (1.07) | 23.60 (1.37) | 21.14 (1.36) | 31.95 (6.90) |
| Completeness (%) | 99.75 (98.28) | 97.44 (83.70) | 99.93 (99.56) | 99.63 (97.83) |
| CC1/2 | 1 (0.504) | 1 (0.509) | 1 (0.65) | 1 (0.971) |
| **Refinement** |  |  |  |  |
| Resolution (Å) | 52.095 - 1.070 | 52.182 - 1.050 | 52.197 - 1.050 | 52.310 - 1.500 |
| Reflections used in refinement | 89185 (8692) | 92346 (7821) | 95045 (9365) | 32953 (3194) |
| Reflections used for R-free | 4517 (451) | 4781 (428) | 4906 (498) | 1608 (150) |
| R-work/R-free (%) | 20.43/21.06 | 19.37/20.21 | 19.91/20.18 | 18.53/20.18 |
| **No. atoms** |  |  |  |  |
| Protein | 1289 | 1290 | 1297 | 1299 |
| Ligand/ion | 18 | 24 | 22 | 31 |
| Water | 120 | 139 | 137 | 140 |
| **B-factors** |  |  |  |  |
| Protein | 16,5 | 14,21 | 15,25 | 15,24 |
| Ligand/ion | 18,83 | 15,04 | 17,56 | 23,09 |
| Water | 23,81 | 21,52 | 22,62 | 22,2 |
| **R.m.s. deviations** |  |  |  |  |
| Bond lengths (Å) | 0,004 | 0,004 | 0,004 | 0,005 |
| Bond angles (°) | 0,85 | 0,76 | 0,75 | 0,79 |

(One crystal for each structure)
\*Values in parentheses are for the highest-resolution shell.

